# Transcriptional regulation of the main olfactory epithelium by environmental olfactory exposures

**DOI:** 10.64898/2026.03.24.713727

**Authors:** Varun Haran, Chinyi Chu, Ryan E. Owens, Thomas J. Mariani, Julian P. Meeks, Regina K. Rowe

## Abstract

The nasal epithelium is a complex tissue composed of both respiratory and olfactory tissue, and is constantly exposed to environmental insults, including toxins and pathogens. The main olfactory epithelium (MOE) serves as the critical site for olfaction, or sense of smell. Dysfunction at this critical barrier tissue can result in partial or total loss of olfactory function, resulting in significant impact to quality of life. The MOE is heterogeneous, comprised of many cell types including olfactory sensory neurons, support cells, and immune cells. It is not well understood how these diverse cell types in the MOE interact to regulate this tissue during homeostasis, and during times of injury and inflammation. We investigated how environmental olfactory exposures impact cell type specific transcriptional responses in the mouse MOE. We performed single-cell RNA sequencing (scRNA-seq) of the MOE following controlled environmental exposure to both well-known odorants and allergens. We identified major cell types and subtypes within the MOE, and identified transcriptional changes in response to the olfactory exposures. We identified transcriptional changes in OSNs, sustentacular cells, and resident immune cells to each condition. This indicated that environmental olfactory exposures drive changes to multiple cell types in the MOE. To our knowledge, this is the first study to identify effects of environmental olfactory exposures on cell-type specific transcription at homeostasis. These findings highlight the potential importance of multi-cellular interactions and communication in regulation of the olfactory epithelium.

## Introduction

The nasal mucosa is constantly exposed to external stimuli, making it a first line of defense against a diversity of non-self insults, including toxins, pollutants, allergens, and pathogens. This tissue also must simultaneously provide the critical function of olfaction as the sole site of odor detection by peripheral olfactory sensory neurons (OSNs). In mammals, olfaction plays a critical role in sensing the external environment to not only protect the organism from danger signals (unpleasant odors)^1^, but is also required for optimal food and nutrition, and social relationships^2–4^. In humans, olfactory dysfunction up to anosmia, which is the complete loss of smell, has been shown to significantly impact quality of life ^5, 6^. Inflammation is one cause of olfactory dysfunction, including postinfectious olfactory dysfunction and chronic rhinosinusitis as two common etiologies ^5, 7^. The observation during the COVID-19 pandemic that anosmia was an early symptom in SARS-CoV-2 infection^8, 9^ highlighted our limited understanding of the complex interactions between olfaction and nasal inflammation. Despite the notable impact of olfactory dysfunction on health and well-being, we still have a limited understanding of how this tissue reconciles its dual roles of sensing inhaled odorants and defending against toxins, pollutants, allergens, and infectious agents.

The main olfactory epithelium (MOE) is highly heterogeneous and is composed of multiple cell types. In addition to OSNs, the MOE includes epithelial, support (sustentacular), solitary chemosensory (microvillus), basal, and resident immune cells ^10–12^. In all sensory epithelial tissues, intercellular interactions between cell types, including neurons and resident immune cells, have been shown to regulate tissue functions and homeostasis ^13, 14^. In the olfactory epithelium, multiple studies describe multicellular interactions to regulate neuronal function and regeneration^12, 15–18^. However, we still have a limited understanding of how external stimulation – either olfactory or toxic – impacts the molecular communication between cells. For example, we do not understand how the MOE maintains OSN function during periods of irritation or inflammation. A better understanding of how chemosensory neurons influence immune cell functions and vice versa during both homeostasis and inflammatory states, could lead to improved treatments of inflammation-induced olfactory dysfunction.

The goal of this study was to determine how environmental olfactory exposures impact cell-type specific transcriptional responses in the mouse MOE. We performed single-cell RNA sequencing (scRNA-seq) following controlled environmental exposure to monomolecular odorants, non-self social odorants and excretions, and allergens. We identified major cell types and subtypes within the MOE, then assessed transcriptional changes in response to each environmental exposure. Our results indicate parallel changes in OSNs, sustentacular cells, and resident immune cells to each condition, providing a foundation for future mechanistic studies into this topic. To our knowledge, this is the first study to identify effects of environmental olfactory exposures on cell-type specific transcription at homeostasis, and the findings highlight the potential importance of multi-cellular interactions and communication in regulation of the olfactory epithelium.

## Materials and Methods

### Mice

Wild-type male and female C57BL/6J mice (Strain #:000664) aged 6 weeks were purchased from the Jackson Laboratory. The mice were kept in home cages with a reverse 12/12 hour light/dark cycle and had unlimited access to food and water. All animal experiments were conducted in accordance with the University Committee on Animal Research of the University of Rochester.

### Environmental exposure conditions

Prior to environmental olfactory exposure, mice were solo housed for 7 days. For soiled bedding exposure, soiled bedding from 8-week-old wild-type male and female BALB/cJ (Strain #000651) mice was collected after 7 days and combined equally. At the start of the experiment (day 0), mice were moved into a new home cage with a petri dish containing 8g of mouse bedding material (1/8” Performance Bedding, Biofresh) spiked with the treatment conditions. For treatments using aqueous solutions, 1mL was mixed with the 8g of mouse bedding. Treatments were as follows: 1) water only (negative control), 2) amyl acetate (0.5%, Sigma Aldrich Catalog #W504009), 3) house dust mite (Dermatophagoides pteronyssinus extract, 2mg/mL; Citeq Biologics, Groningen, Netherlands, Catalog #15J01); or 4) 8g of soiled bedding material. The mice underwent a cage change every two days (days 0, 2, 4, 6), accompanied with fresh stimulus material. One male and one female mouse were used for each treatment condition per experiment. The experiment was repeated in duplicate for a total of 4 mice (2 male, 2 female) per exposure condition in the final data set. An overview of this experimental design is shown in Figure 1.

**Figure 1.**
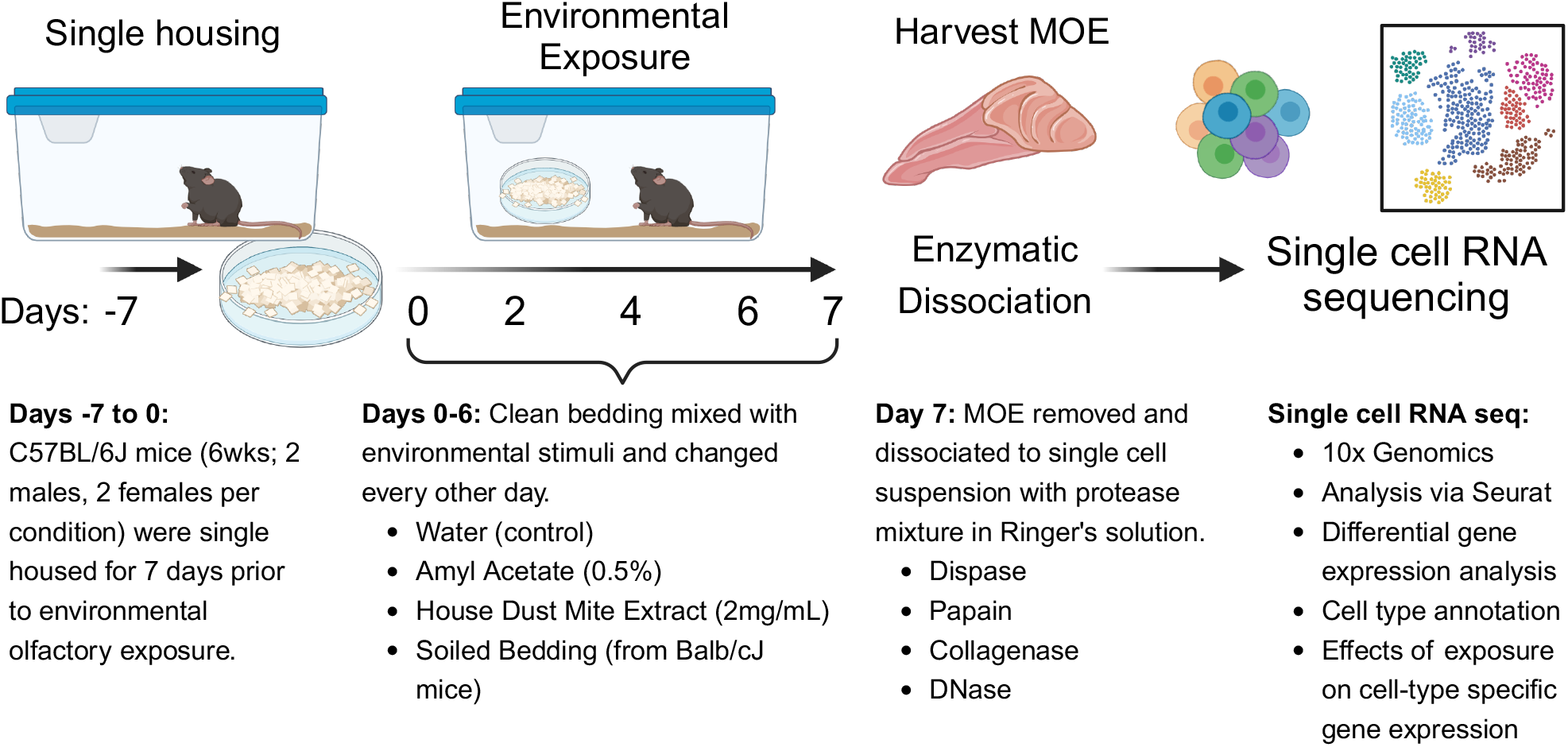
Overview of Environmental Olfactory Exposure. Mice were single housed for 7 days prior to environmental olfactory exposure. Environmental stimuli were applied to clean bedding for 6 days. The main olfactory epithelium was harvested, dissociated, and single cell RNA sequencing performed.

### Main olfactory epithelium dissociation procedure

At day 7, mice were euthanized by isoflurane inhalation and decapitation. The head was hemisected sagittally along the septum, exposing the turbinates with the olfactory epithelium. The turbinates were then removed from the nasal cavity and collected in freshly prepared oxygenated Ringer’s solution containing 115 mM NaCl, 5 mM KCl, 2 mM MgCl2 hexahydrate, 2 mM CaCl2 dihydrate, 25 mM NaHCO3, 10 mM HEPES, and 10 mM glucose with a pH of 7.4 at 4°C. The olfactory epithelium was carefully dissected from the turbinates and minced with fine forceps under a dissection microscope (Leica Microsystems, Buffalo Grove, IL, USA). This cell suspension was then transferred to a cocktail of tissue digestive enzymes containing Dispase II (2 mg/mL, Sigma Aldrich Catalog #D4693), Collagenase Type 1 (2 mg/mL, Sigma Aldrich Catalog #SCR103), Papain (0.25 mg/mL, Sigma Aldrich Catalog #10108014001), and DNase (2 mg/mL, Sigma Aldrich Catalog #DN25) in Ringer’s solution for 45 minutes in an orbital shaker incubator at 37°C. The reaction was then terminated with Ringer’s-FBS (FBS-10%) and the cells were triturated, filtered, and centrifuged. The cell suspension was filtered once more with a cell strainer with 40-micrometer diameter pores and then subjected to centrifugation. The final cell suspension was resuspended in 500 μl of Ringer’s solution and 10 μl was set aside for cell viability assessment by Trypan blue staining.

### Single cell RNA sequencing (scRNAseq) of mouse olfactory epithelial cells

Post-dissociation, single cell suspensions were used for single cell capture via 10x Genomics platform by the University of Rochester Genomics Resource Core. Samples were first barcoded using the 10x Genomics CellPlex cell hashing protocol per manufacturer’s recommendations (10x Genomics, Pleasanton, CA). Barcoded samples were pooled and a goal of 10,000 cells were captured on the 10x Genomics Chromium platform. Pooled samples from 4 MOEs (one male or female from each exposure condition) were captured across 2 independent captures to facilitate sufficient cells per animal/sample and decrease batch effects. Total RNA was isolated, cDNA libraries constructed, and libraries were sequenced on the Illumina NovaSeq6000 (Illumina, San Diego, CA). Tissue harvesting, dissociations, and captures were repeated on subsequent days to perform the experiment on 8 total animals (1 male and 1 female for 4 conditions) in a sequencing experiment. This experiment was performed twice for a total of 16 animals (8 male, 8 female, 4 per condition) for a total of 50,032 cells in the final analyzed data set.

### Data Integration, Clustering and Cell Annotation

Quality control, cell subset analyses, and differential gene expression were performed using Seurat (version 5.3^19^). Histograms of feature numbers were generated to identify filtering criteria. Cells with more than 8,000 or less than 200 features and those exceeding a mitochondrial percentage of 20% were excluded in the analysis. The mitochondria gene percentage is slightly higher than reported by prior publications^10, 20, 21^, potentially due to the environmental exposure protocol. The data were normalized using the SCTransform method and data integration occurred through RPCA. Differentially expressed genes were identified and the top 2,000 genes were used to perform PCA analysis. PCA analysis was then used to identify clusters and UMAP transformation was performed. Subclusters were identified similarly by performing clustering of each main cluster. Marker genes of each cluster were identified using the FindAllMarkers function via Wilcoxon rank-sum test. Cell type assignments were performed using a combination of methods: 1) marker genes from individual clusters were used to determine the cellular characteristics using Azimuth database (Supp. Table 1), and 2) expression levels across clusters of well-known and canonical cell-type specific genes were used based on previous studies^12, 21^.

### Differential Gene Expression Analysis

Differential gene expression analysis was performed using MAST (Model-based Analysis of Single-cell Transcriptomics) method to identify genes that were different between exposure groups when compared to control group (water) within each cell cluster using Seurat FindMarkers function. Genes that were infrequently detected were removed and only genes that were detected in at least 10 percent of cells in either of the exposure groups or control (water) group were used in differential analyses. Genes were identified as significantly different if adjusted p-value is less than 0.05 and the estimated fold-change was greater than 1.5-fold (Supp. Table 2).

### Pathway Enrichment Analysis

Genes identified as differentially expressed in within each cluster in exposure groups, compared to control were subsequently used for canonical pathway enrichment identification through Enrichr (https://maayanlab.cloud/Enrichr/) utilizing Mouse Gene Atlas, PanglaoDB_Augmented_2021, and Azimuth_Cell_Types_2021. Canonical pathways were reported when the p-value was less than 0.05 (Supp. Table 3). The top 10 pathways for each cluster were identified based on a combination of Enrichment Score and p value<0.001. Graphical representation was performed for these top 10 pathways in R using ggplot2 package.

## Results

### Cell type transcriptional diversity in the olfactory epithelium

Using a mouse model, we designed a method to administer environmental olfactory stimulation followed by isolation of the MOE to determine how olfactory stimulation in the environment impacts MOE cellular transcriptional responses. A diagram of our approach is shown in Figure 1. Mice (C57Bl6/J) were single-housed for 1 week prior to stimulation to minimize exposure to non-self and other uncontrolled environmental odorants. Mice were then provided odorant stimuli, applied to fresh bedding, in a petri dish. Stimuli chosen were water (control/diluent), amyl acetate (0.5% in water), soiled bedding from male and female Balb/cJ mice, and house dust mite extract (1 mg/mL in water). Fresh stimuli were provided every-other-day for 6 days, with MOEs harvested one day after the last odorant exposure (Day 7). Tissue was dissociated and single cell RNA sequencing (scRNAseq) performed (Fig. 1). This experimental design provided environmental exposure to stimuli in a naturalistic context and included stimuli representing different olfactory experiences: amyl acetate (monomolecular odorant), soiled bedding (a non-self, natural odorant mixture), and house dust mite extract (a foreign environmental mixture with known immunomodulatory components).

Single cell RNA sequencing was performed on dissociated MOEs and cell type specific annotations were identified. UMAP projections of cells within 15 major clusters and are shown in Figure 2. These clusters corresponded to major cell populations known to be present in the olfactory epithelium and adjacent respiratory epithelium (Fig. 2B). Using subsequent sub-clustering methods, we identified additional subsets for many of the larger clusters (Fig. 2A-B, Supp. Fig. 1). Although several single cell RNA sequencing-based datasets exist for the olfactory epithelium^10–12, 22^, we did not assume that our procedures isolated the same tissue areas, or the same gene expression patterns as previous studies. Instead, we assigned cell types to clusters based on two approaches. First, we used the Azimuth database, which produces a ranked list of cell type assignments based on assignments from other studies. Secondly, we manually compared marker gene expression (Supp. Table 1) patterns for each cluster and subcluster to existing data sets ^12, 22^, including non-RNA-sequencing-based studies (Fig. 2, Supp Fig. 1, Supp. Fig. 2).

**Figure 2).**
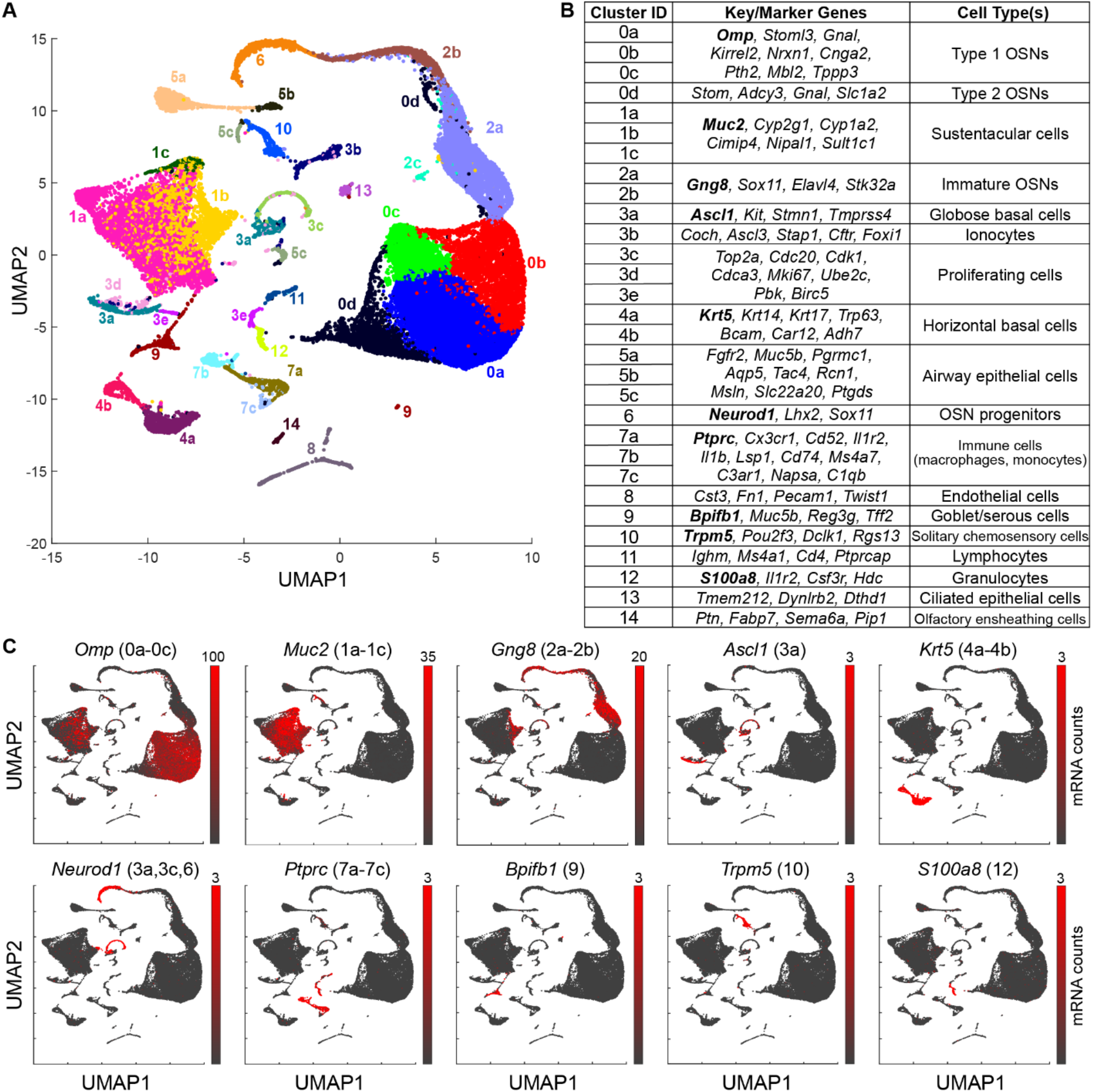
Cell type inferences from single-cell RNA sequencing of the dissociated main olfactory epithelium (MOE). **(A)** Uniform manifold approximation and projection (UMAP) plot of transcriptional patterns taken from 50,032 MOE cells from 16 mice (4 experimental conditions, 2 males and 2 females per condition). Initial clustering identified 14 clusters, some of which were subclustered (see Figure S1), resulting in a total of 31 clusters, noted by labels and unique colors in the plot. **(B)** List of clusters (Cluster ID) and associated genes expressed that assisted in cell type identification. Note: Cluster 2c is not listed to conserve space. **(C)** Colorized expression plot of raw mapped mRNA counts for key/marker genes for a subset of clusters/cell types. The max of the colorization range (pure red hue) reflects the 95^th^ percentile expression level for genes expressed in the dominant cell populations, the 99.9^th^ percentile of expression (for genes expressed in rarer cell populations, minimum 3 counts). Additional marker genes are plotted in Figure S1.

The largest resulting major cluster (Cluster ‘0’) included mature olfactory sensory neurons, consistent with procedures effectively isolating and dissociating the MOE. Within Cluster 0 we identified 4 distinct subclusters: Subclusters ‘0a-0c’ expressed genes consistent with type 1 olfactory neurons expressing marker genes *Omp, GnaI, Cnga2*, while Subcluster ‘0d’ expressed *Stom, Adcy3, and GnaI*, markers of type 2 olfactory neurons. Cluster 1 expressed markers of sustentacular cells including *Cyp2g1, Cyp1a2*, and *Muc2*, and we identified 3 subclusters in this group. Cluster 2 expressed genes associated with immature olfactory sensory neurons with two distinct subclusters. Cluster 3, which initial cluster analysis suggested had the 4^th^ most cells overall, was highly heterogenous. Subclustering readily identified the presence of 5 separable subsets (Subclusters ‘3a-3e’), including globose basal cells (‘C3a’) and ionocytes (‘C3b’). The identification of these clearly separable cell types within a single major cluster highlighted the importance of our subclustering approach in this complex tissue. Other noteworthy cell types included horizontal basal cells (‘C4’), OSN progenitors (‘C6’), other proliferating cells (‘C3’), endothelial cells (‘C8’), solitary chemosensory cells (‘C10’), and olfactory ensheathing cells (‘C14’). Airway and respiratory epithelial cells were identified across 3 clusters (‘C5’, ‘C9’, and ‘C13’). Cluster 5 had overlapping genes for airway epithelial and sustentacular cell markers, with Cluster 9 and 13 consistent with goblet/serous and ciliated cells, respectively.

Immune cell populations were distributed across multiple clusters, consistent with the anticipated diversity and proportions of specific immune cell types. The largest cluster was Cluster 7, highly expressing the common leukocyte antigen gene, *Ptprc*, and markers of myeloid cell origins (monocytes and macrophages), which included *Cx3cr1, Cd52, Ms4a7, Il1r2, and Il1b*. This subcluster could be separated into three distinct cell types as demonstrated by differential expression of *Cx3cr1*, a marker typically expressed in tissue resident macrophages (‘C7a’). In contrast, Subclusters ‘7b’ and ‘7c’ were more consistent with dendritic cells, monocytes, and monocyte-derived macrophages. While the myeloid population was the most prevalent, additional clusters were identified with marker genes for B and T lymphocytes (‘C11’; B cells: *Ighm, Ms4a*; T cells: *Cd4, Cd3d, Cd3e*), and granulocytes (‘C12’; neutrophils: *Il1r2, Csf3r)*. Based on this analysis, 31 total cell types were identified, reflecting the diversity of cell types in this sensory tissue.

### Effects of environmental olfactory exposure on transcriptional responses

To investigate transcriptional changes in MOE cell types following environmental exposures, we performed differential gene expression analysis. We first determined the distribution of each exposure condition with respect to the total cells in the data set as shown in Figure 3. We confirmed that each exposure condition did contain cells across every cluster and subcluster. We then tested for overall proportional differences across clusters for each condition. Chi-squared analysis did detect differences in proportions when all conditions were considered (Fig. 3B). For example, the OSN cluster, C00a, had a smaller proportion of cells from the house dust mite exposed condition. In contrast, this condition increased proportion in the main sustentacular cell cluster, C01a, and innate cell immune clusters C07 (monocytes/macrophages) and C12 (neutrophils) (Fig. 3B), potentially indicating tissue remodeling as a result of the inflammatory stimulus. As expected, smaller clusters were more impacted by variation across conditions, *e*.*g*. C11-C14, which may simply be due to overall lower cell numbers in these clusters. However, no new cluster or subcluster was specific to a given exposure. Despite these proportional variations, each cluster contained cells from every treatment condition (Fig. 3C), allowing for subsequent comparisons of transcriptional responses across exposures.

**Figure 3).**
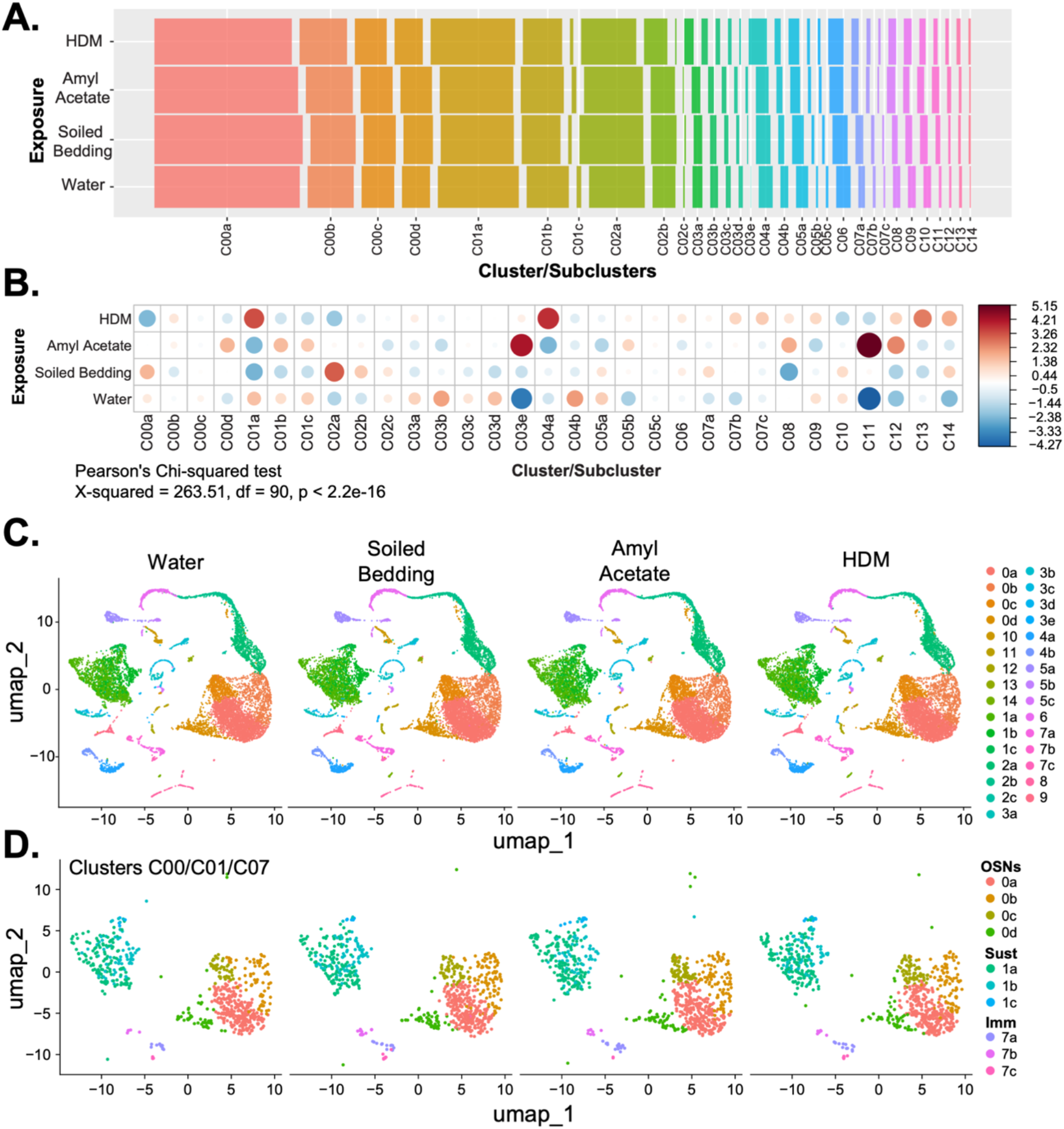
Distribution of cells across clusters following environmental olfactory exposures. The proportion of cells in the single cell RNA sequencing data set were determined for each exposure condition: water, soiled bedding, amyl acetate, and house dust mite (HDM). (A) Bar graph of the number of cells in each cluster and subcluster for each exposure condition. (B) Chi-squared analysis of data in (A) demonstrating differences in proportion across clusters and subclusters for each condition. Dot plot of Pearson’s residuals with size of dot representing the size of contribution to the χ^2^ value, color represents direction of association (red = positive; blue = negative). (C) UMAP plot demonstrating that every cluster is represented in each exposure condition with (D) highlighting clusters of interest used in subsequent pathway analysis (C00, OSNs; C01, sustentacular cells; C07, immune cells).

Using the water-only stimulus as the control, we evaluated differential gene expression for each exposure condition. Differentially expressed genes were identified for all exposure conditions, including both upregulated and downregulated genes (Fig. 4A). The number of differentially expressed genes varied dramatically by cell type, and varied with sex (Fig. 4A). Sustentacular cells (‘C01a-b’), immature OSNs (‘C02a’), horizontal basal cells (‘C04a’), immune cells (‘C07a-b’), solitary chemosensory cells (‘C10’) and olfactory ensheathing cells (‘C14’) showed evidence of stronger shifts in their gene expression patterns. In some cases, the number of differentially expressed genes in a cell type varied by sex (Fig. 4A). Although sex differences were apparent in the number of differentially expressed genes in some clusters (for example, more upregulated genes in female sustentacular cells ‘C01a-b’ than in males, or more downregulated genes in immune cell ‘C07a’ in males than in females), it is important to note that male and female samples (from all stimulus conditions) were pooled in males separately from females. This could lead to subtle batch effects that persisted despite computational integration steps designed to minimize such effects.

**Figure 4).**
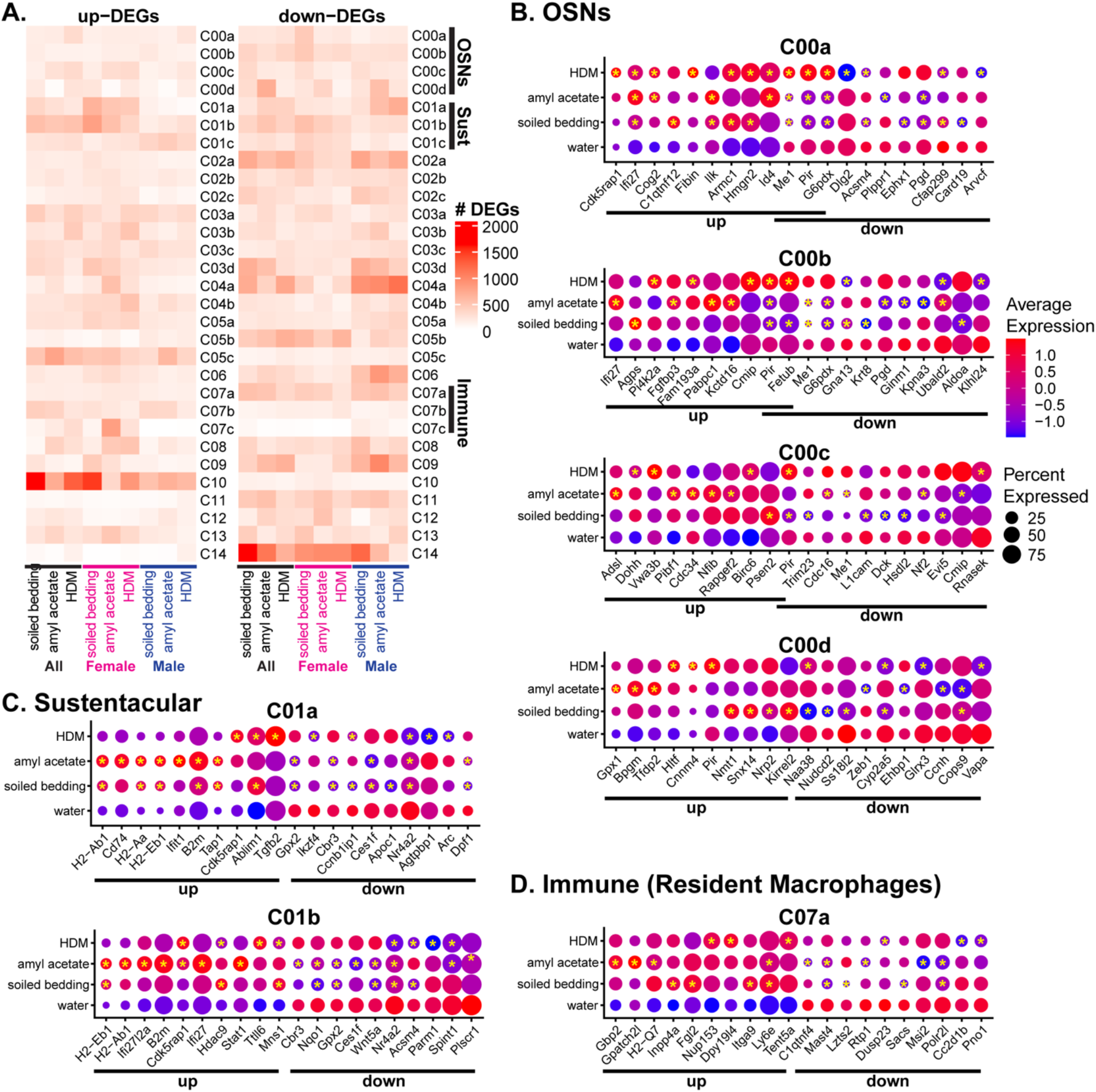
Environmental olfactory exposure results in differential gene expression across multiple cell clusters. Differential gene expression analysis was performed via Wilcoxon rank testing for each exposure condition versus the water (A) control: BA (soiled bedding/water), CA (amyl acetate/water), DA (HDM/water). Comparisons were also performed for each sex (male and female mice) for each exposure. (A) heatmap of the number of differentially expressed genes for each cluster and subcluster. (B) Dot plots of the top DEGs (up and down) for: OSN (C00), Sustentacular (C01), and Immune (C07) subclusters. Gene expression is shown for each exposure condition and were either up or downregulated as compared to water for at least 1 treatment. Top genes were determined by p value <0.01 and log2fold change. Significant comparisons versus water are noted by * on each dot.

To more closely investigate gene expression changes, we performed targeted analysis of selected genes from the OSN (‘C00’), sustentacular (‘C01a-b’), and immune cell (‘C07a’) clusters. The top differentially expressed genes for each subcluster are shown in Figure 4B-D, with each assessed gene either significantly up- or down-regulated for at least one treatment compared to the water control. In some cases, top differentially expressed genes had well-established functions in the cell types expressing them. For example, the genes *Nrp2* and *Kirrel2* are both associated with axon targeting and neural circuit formation in OSNs and were up-regulated in response to soiled bedding.(Fig. 4B, ‘C00d’). In others, top differentially expressed genes had well-established functions, but were not strongly associated with the cell types expressing them (*e*.*g*., upregulation of major histocompatibility genes *H2-Ab1, H2-Aa*, and others in sustentacular cells; Fig. 4C, ‘C01a’ and ‘C01b’). Finally, some genes reflected tissue-specific specialization such as up-regulation of *Inpp4a* may suggest and anti-inflammatory role during homeostasis (Fig. 4D, ‘C07a’). Each stimulus was associated with multiple up- and down-regulated genes in these cell types, producing a complex ensemble of transcriptional differences (Fig. 4).

To help interpret the complex differential gene expression patterns we observed, we turned to pathway analysis using Enrichr databases, again comparing each stimulus condition to the control (water). Numerous pathways were identified (Figure 5; Supp. Table 3), and while there were differences in overall pathways associated with a given stimulus condition, common themes were evident in specific clusters and cell subsets (Figure 5). In mature OSNs (Cluster 0), pathways of both up- and down-regulated genes following olfactory stimulation were largely related to cellular responses to stress, apoptosis and cell death, mitochondrial and metabolic processes, protein translation, and neuronal/axon growth. Pathways of up-regulated genes in sustentacular cells (Cluster 1), were largely related to antigen processing and presentation, and immune and inflammatory responses, while downregulated gene pathways represented metabolic, stress response, and protein regulation pathways were downregulated. In immune cells (Cluster 7), including macrophages and monocytes, pathway analysis differed across subsets. Downregulated gene pathways in C07a, which corresponds to the resident macrophage population, were related to RNA and protein regulation processes such as RNA splicing, ubiquitination, and autophagy. Upregulated gene pathways in C07a were associated with anti-viral and anti-bacterial immune responses, and pro-inflammatory processes. In contrast, circulating or non-resident monocytes (Cluster C07b) had downregulation of genes related to core cellular processes including ribosomal regulation and protein initiation and translation (Fig. 5, Supp Table 3).

**Figure 5:**
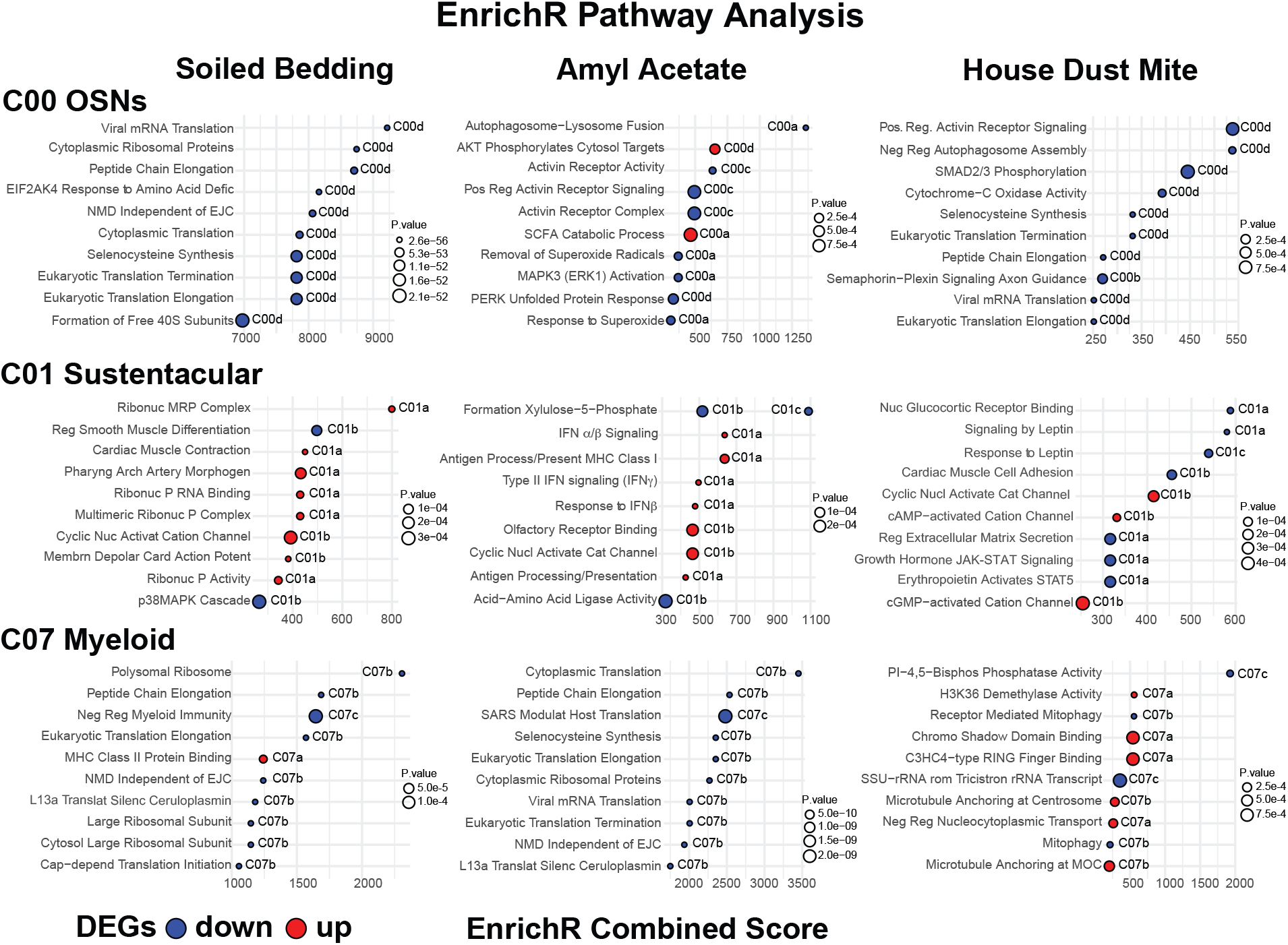
Pathway analysis was performed using EnrichR (GO Biological, GO Cellular, GO Molecular, Reactome, KEGG, and WikiPathways) using the differentially expressed genes identified for each treatment as compared to water control. Shown is the combined EnrichR score for the top 10 pathways for each cluster (C00/OSN; C01/Sust, C07/Immune) and condition (p<0.001). Dot color represents either down (blue) or up (red) regulated genes, and size of the dot is p value.

## Discussion

This study evaluated the effects of specific types of environmental stimulation on the diverse cell types found in the mouse MOE. We did so while establishing an experimental pipeline that involved stimulating the MOE with controlled environmental substances, followed by dissociation, cell indexing, single cell RNA sequencing, and integrated gene expression analysis. This is one of relatively few studies to date to independently characterize the diversity of cell types in this heterogeneous tissue ^10–12, 22^. Our results show that in response to environmental olfactory stimulation, multiple cell types in the OE, including neurons, sustentacular, and immune cells, undergo stimulus-dependent transcriptional changes (Figs. 3–5). This experimental paradigm includes the capacity to investigate sex-specific effects of these responses, although this specific study was not optimized for such comparisons. To our knowledge, this is the first study to identify transcriptional changes in the mouse MOE in response to environmental stimulation in multiple cell types, including immune cells.

The motivation for this study, and the development of this experimental platform, was to investigate the mechanisms by which the MOE maintains its function in the constant presence of harmful or helpful stimuli. Using single cell RNA sequencing, we sought to identify how the diverse cell types within the MOE respond to environmental olfactory stimuli. Although single cell RNA sequencing can only provide a limited snapshot of the overall changes experienced by the MOE, it does so in a way that supports objective, systematic comparisons across cell types. Here, we identified transcriptional changes specific to each stimulus, including some that were consistent with the different categories of exposures chosen. For example, one of the chosen stimuli was soiled bedding from a mouse strain (BALBc\J) different than the C57Bl6\J mice being studied. Mice of different strains emit highly complex mixtures of non-self chemosensory cues in their dander, tears, urine, and feces ^23–25^. We and others have shown that urine and feces stimulate specific sets of chemosensory neurons in a ligand- and concentration-dependent fashion^26–29^. Given the complexity of these unfamiliar non-self chemosensory cues, we expected that this exposure would drive widespread transcriptional changes in OSNs (Cluster ‘C00’). In contrast, amyl acetate, a well-characterized monomolecular odorant, would be expected to activate only a small subset of OSNs selectively expressing one of about a dozen known amyl acetate-sensitive olfactory receptors ^30^. Even for this limited environmental stimulus, we observed transcriptional changes in OSNs compared to the negative control water stimulus.

Intriguingly, we also identified differential gene expression patterns to these “pure olfactory” stimuli in other cell types, including sustentacular (C01) and immune (C07) cells. In sustentacular cells, differentially expressed genes were associated with many cellular processes consistent with these cells’ barrier and support functions, including cyclic nucleotide-gated ion channel activity, epithelial/barrier function, and metabolism. However, many upregulated genes in the sustentacular cell clusters were also related to immune responses and signaling, including antigen presentation and cytokine responses. This suggests a critical role for sustentacular cells in maintaining epithelial integrity, supporting olfaction, and regulating local inflammatory responses.

Many differentially expressed genes were noted to have overlap across treatment conditions, with more overlap noted between soiled bedding and amyl acetate (data not shown) as compared to house dust mite exposure. This would suggest that activation of the OSNs in response to these non-toxic olfactory stimuli triggers similar responses across the surrounding cellular milieu. While the response of the tissue to a more inflammatory trigger, house dust mite, results in divergent responses. Pathway analysis supported the involvement of many pathways involved in stress responses, cell death, and metabolism – cellular responses which would need tight regulation to maintain tissue function and homeostasis.

Resident immune cells are poised to respond quickly to inflammatory insults, including microbes and toxins. Compared to the predominant cell types in the MOE (OSNs, sustentacular cells, microvillous cells, etc.), our understanding of the role of resident immune cells in this tissue is limited. These cells are likely involved in neuronal repair and regeneration in the setting of inflammation or tissue damage ^16, 31–33^. Neurons in other epithelial tissues, such as the skin and gut, are known to be regulated by resident immune cell activity ^34, 35^. We hypothesized that this also occurs in the MOE, and that neuroimmune interactions could be specifically regulated to preserve olfactory function during inflammatory exposures.

Here, our methods supported the advancement of our understanding of the diversity of resident immune cells in the MOE, both at homeostasis and in the presence of olfactory and inflammatory stimuli. Immune cells make up a small proportion of the total MOE tissue, about 4-6% (Fig. 2). Because of their relatively low abundance, we were only able to clearly identify major immune cell subsets, including myeloid (macrophages, monocytes, dendritic cells; C07), lymphocytes (B and T cells; C11), and granulocytes (neutrophils, eosinophils; C12). Resident macrophages (C07a), which are positive for the canonical marker *Cx3cr1*, represented the largest immune cell population. Microglia in the brain also express high levels of *Cx3Cr1* ^36^. Our data supports that resident macrophages in the MOE play a similar role to that of microglia in the brain, potentially supporting pro-inflammatory and anti-inflammatory functions ^36^. Even in response to “pure olfactory” stimulation, many differential gene expression pathways were associated with phagocytic, antigen presentation, and pro-inflammatory responses.

In addition to the effects we saw in non-neuronal cells following environmental olfactory stimulation, we also saw evidence of differential gene expression in OSNs (C00) following environmental stimulation with house dust mite extract, a well-established source of immunostimulatory molecules, including house dust mite antigen and lipopolysaccharide^37^. This environmental exposure paradigm, which introduced mice to this substance consistently, but non-invasively, produced relatively mild changes in pro-inflammatory gene expression in immune cells and sustentacular cells (Figs. 4–5). It is not clear whether these gene expression changes reflect altered chemosensory function (*e*.*g*., changes in sensitivity to odorants, adaptation to repeated odorant exposure, etc.), which will require future studies to further investigate.

Although this approach cannot test whether these changes reflect direct communication between MOE cell types, these results, taken as a whole, indicate that environmental drivers of OSN activity have the capacity to alter the sustentacular and resident immune cell function. Further studies will be needed to test hypotheses relating OSN activation to the integrity and function of the olfactory epithelial barrier by sustentacular cells, and local inflammatory responses to toxins, allergens, and pathogens by resident immune cells.

## Funding Sources

University Research Award from the University of Rochester Medical Center; the University of Rochester Department of Pediatrics; T32ES007026 (RO) from NIEHS/NIH; and partial support provided by R01DC017985 (JPM) from the NIDCD/NIH, and K08AI163380 (RKR) from the NIAID/NIH.

## Acknowledgements

We want to thank Bailey Matthews and Michael Mastrangelo for their technical assistance. We acknowledge the staff of the University of Rochester Medical Center Genomics Resource Core for assistance with single cell RNA sequencing data acquisition.

---

**Figure S1).**
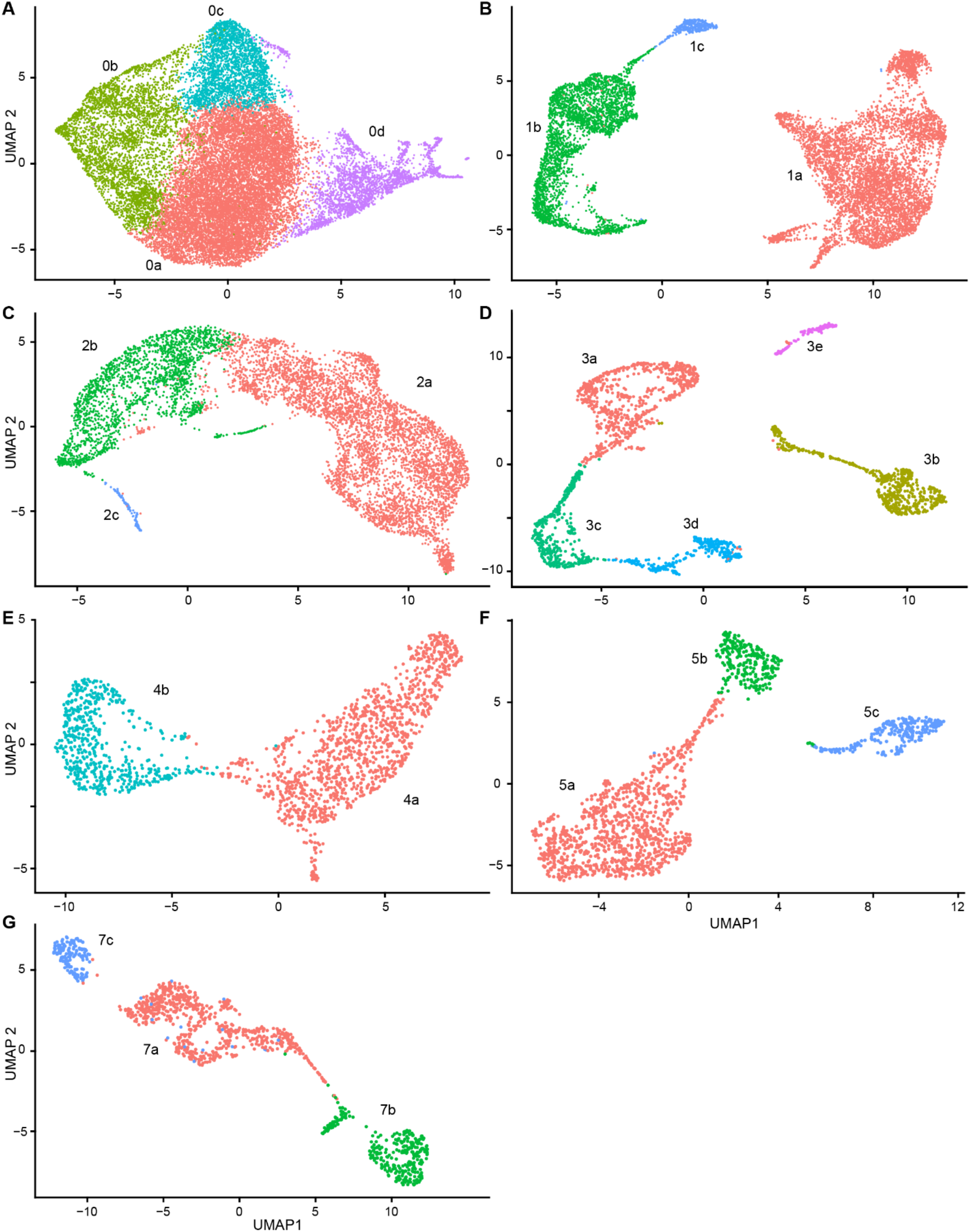
UMAP plots of subclustering from Figure 2. (**A-G**) Colorized maps for clusters 0, 1, 2, 3, 4, 5, and 7, respectively. Parameters for clustering were adjusted based on the sample sizes, with lower thresholds for cluster separation (“resolution” parameter in Seurat::FindClusters) used for clusters with smaller overall numbers of cells (range 0.08 to 0.025).

**Figure S2).**
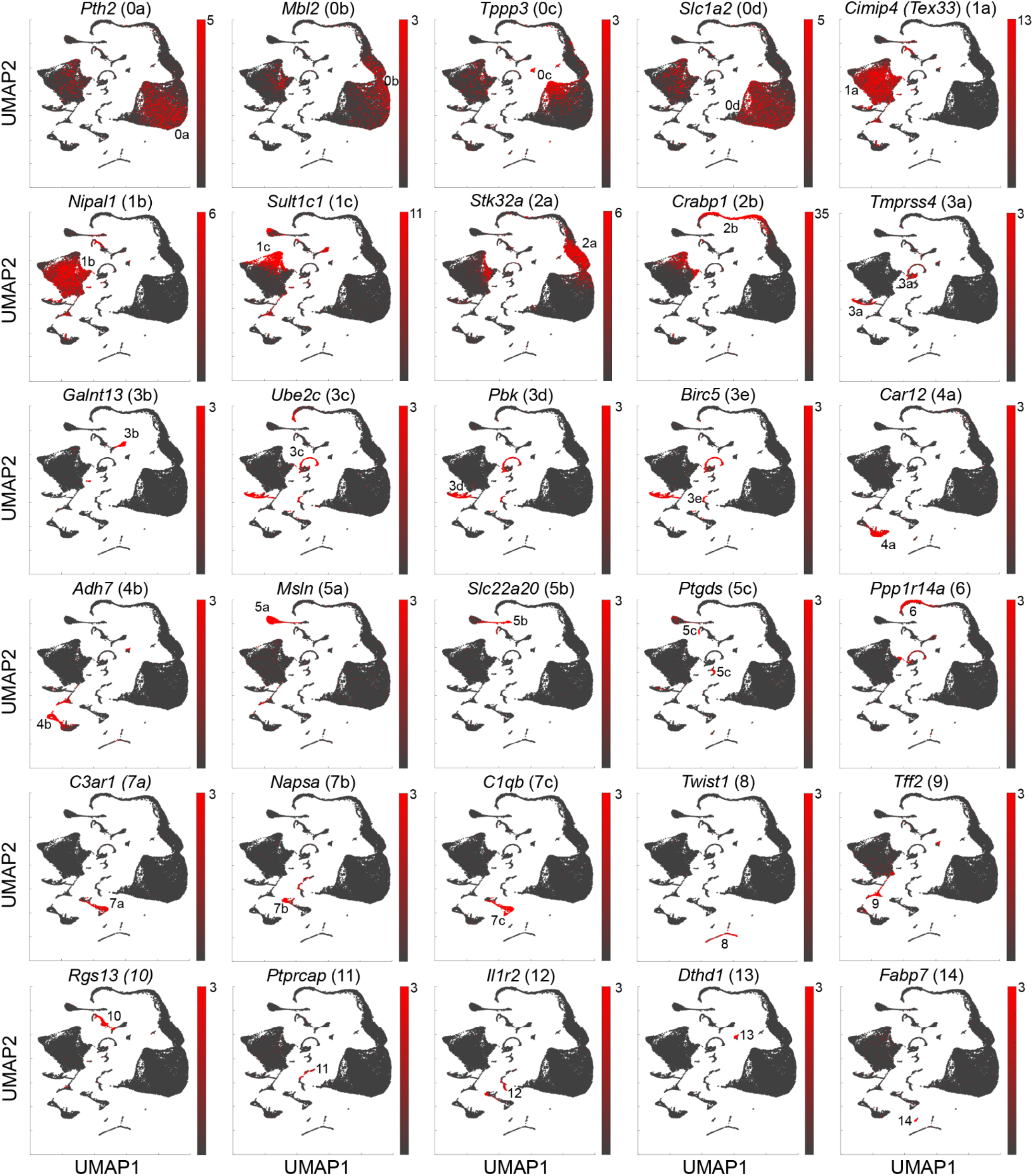
UMAP heat plot of enriched genes for each cluster. Genes chosen reflect those with highest “selective expression index (See Methods) that were not plotted in Figure 2.

## Notes

### Competing Interest Statement

The authors have declared no competing interest.

